# In Silico Target Identification Highlights IL12B as a Candidate for Small Molecule Drug Development

**DOI:** 10.1101/2024.09.03.609261

**Authors:** Trevor J Tanner

## Abstract

Drug discovery’s rising costs and complexities require innovative strategies to identify viable therapeutic targets. We developed a computational pipeline to pinpoint protein targets lacking known small molecule probes, focusing on sites traditionally considered challenging for small molecule intervention but validated by FDA-approved biologics. Our approach integrates machine learning, public databases, structural modeling, and functional annotations to prioritize novel binding pockets that overlap biologically validated interfaces. This method identified IL12B as a promising candidate, revealing a previously unexploited surface pocket that overlaps part of the briakinumab epitope. Static protein solvent mapping and dynamic fragment simulations provide convergent evidence of druggability, including fragment-binding clusters and chemically diverse hotspots. While not yet experimentally validated, this site represents a plausible target for orally available IL12B inhibitors. Such compounds could address current clinical limitations of antibody therapies - such as prolonged systemic exposure and infection risk - by enabling a shorter half-life and improved mucosal penetration in diseases like inflammatory bowel disease.

## Introduction

The discovery of new medical drugs has become increasingly challenging and costly despite significant technological advancements (Scannell et al., 2012). As the easier targets have been exhausted, scientific progress now requires tackling more complex challenges.

Our research strategy addresses these challenges by leveraging existing knowledge to uncover novel opportunities. Inspired by Sir James Black’s insight that “the most fruitful basis of a new drug is to start with an old drug” (Raju, 2000), we hypothesized that protein sites traditionally deemed “difficult-to-drug” for small molecules—but already validated as therapeutic targets by FDA-approved biologics—may hold untapped potential. Small molecule drugs possess distinct pharmacokinetic and pharmacodynamic properties, such as shorter half-lives and enhanced tissue penetration, which could offer clinical advantages over biologics in certain contexts (Wan, 2016).

This study introduces a computational pipeline designed to identify protein sites lacking known small molecule binders but overlapping—at least partially—with biologic binding epitopes. Our approach builds on the proven efficacy of the SiteMap tool in identifying challenging protein sites, further refining its outputs through machine learning and filtering criteria intended to reduce the risk of failure in downstream campaigns (Liu & Altman, 2014).

To evaluate potential druggability within these sites, we combined two complementary fragment mapping tools: FTMap and SILCS-Hotspots. While FTMap has demonstrated a high true positive rate in identifying druggable regions, its rigid-body framework does not account for protein flexibility. Conversely, SILCS-Hotspots incorporates protein dynamics but tends to overpredict, leading to a higher false positive rate (Ghanakota & Carlson, 2016). By integrating both approaches, we aim to balance structural precision with conformational breadth, enabling more robust identification of functionally meaningful binding sites suitable for future small molecule targeting.

## Methods

### Homologous count file creation

We implemented an initial sequence-based filtering step to reduce the likelihood of off-target binding during drug-target selection. This computationally inexpensive approach prioritized candidate proteins that exhibited minimal primary sequence homology to other proteins in the human proteome. However, this method does not account for proteins that share structurally similar binding sites despite lacking primary sequence homology, a phenomenon that can still mediate significant off-target interactions (Duran-Frigola et al., 2017).

We created a custom bioinformatics pipeline using Nextflow, “bioinfo.nf”, to filter out drug targets with significant primary homology to other human proteome proteins (Di Tommaso et al., 2017). We used AlphaFold structural predictions (version one) of the human proteome as input (Jumper et al., 2021). We extracted FASTA protein sequences from the input PDB files and blasted them against the UniProt reference human proteome using blastp (Altschul et al., 1990). Hits were filtered using a custom Python script, “filter_blast.py”, with a 1E-62 maximal e-value cutoff (Louie et al., 2009), then re-aligned to the query protein sequence using mafft (Katoh & Standley, 2013). Hit counts for each query were tabulated using a custom bash script, “extract.sh”, and saved to a reference file. Fig. 1b visualizes the process.

**Figure 1.**
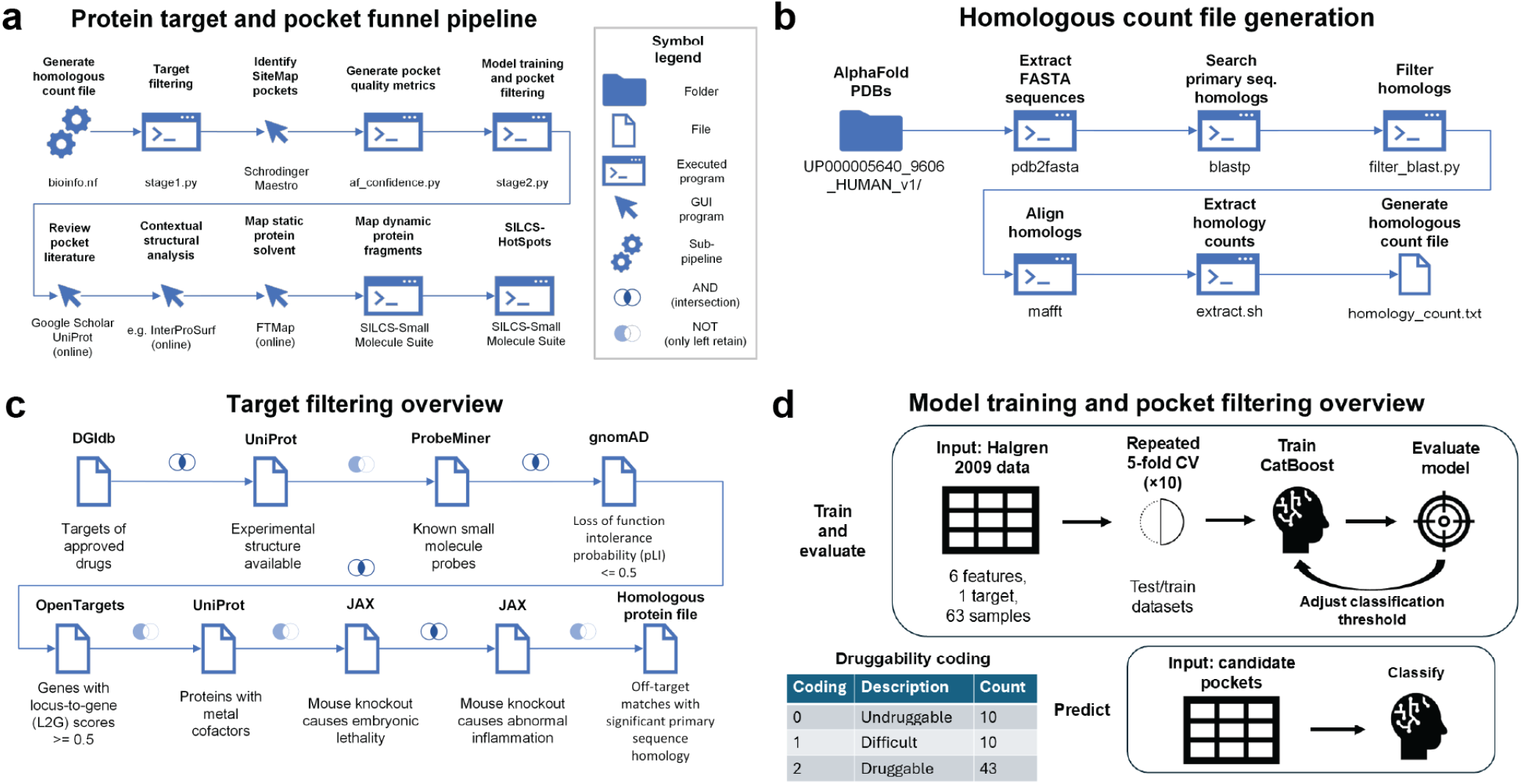
Target evaluation and binding pocket prediction. (a) All intermediate steps and interactions with graphical user interface (GUI) based programs are performed manually. (b) All steps are executed using the Nextflow automation tool. Command arguments not listed. (c) All flow chart steps contained within Stage1.py. Further descriptions of databases and justifications for filtering steps found in methods. (d) All machine learning steps within Stage2.py. Target threshold refers to minimal value for differentiating between druggable and undruggable pockets. CV stands for cross validation.

### Initial target filtering

We applied a series of filters using external and internal data sources to limit the number of protein candidates for structural analysis in a script titled “stage1.py”. We used the Drug Gene Interaction Database (DGIdb) to identify approved drug putative targets (Cannon et al., 2024). We then filtered targets to include only those with experimental structures available in the UniProt database (The UniProt Consortium et al., 2025).

We applied an antibody structure filter using the Thera-SAbDab database to include only targets where a structure is available with a biologic binding (Raybould et al., 2020). This ensured that actual biologic binding sites were characterized and known for our analysis.

We excluded targets with existing small molecule probes available according to the ProbeMiner database to focus on unexplored targets and maximize potential for novel discoveries (Antolin et al., 2018). In the final stage of initial filtering, we considered targets only if they had no significant primary sequence homology to other proteins in the human genome to reduce the probability of off-target effects. The process of initial target filtering is visualized in Fig 1b.

### SiteMap pocket identification

After initial target filtering, we manually loaded structural predictions from AlphaFold into the Schrodinger Maestro application and processed them with the Protein Preparation Wizard using default parameters and the OPLS4 force field (Madhavi Sastry et al., 2013; Lu et al., 2021). We then detected protein pockets using SiteMap with default parameters and exported them to a comma-separated file (Halgren, 2007).

### Pocket quality metric generation

We processed the SiteMap pocket predictions in a custom Python script, “af_confidence.py”, that calculated the percentage of “very high” (>90 pLDDT) quality amino acid predictions in the pocket and the percentage of “very high” quality amino acids in the entire region from the first amino acid in the pocket to the last amino acid in the pocket. The calculated metrics were added as new columns to the original SiteMap dataset and saved to a new comma-separated file.

### Model training

Training data points for Schrödinger SiteMap pockets and druggability labels were extracted from prior research (Halgren, 2009). The features used for model training included SiteMap’s core pocket descriptors, such as Dscore, SScore, size, enclosure, philic, and phobic values, which numerically characterize pocket geometry and physicochemical environment. Druggability labels of “undruggable,” “difficult,” and “druggable” were coded as 0, 1, and 2, respectively.

A CatBoost classifier was utilized as the supervised classification model, selected because its probability outputs can be directly interpreted as a measure of model confidence (Prokhorenkova et al., 2019). Model parameters included 1000 iterations, a learning rate of 0.05, a depth of 6, and a logloss function, with class weights set dynamically based on class imbalance observed during training. Random initialization state was fixed at 42.

Druggability labels were converted into binary values, marking labels as “true” if they exceeded thresholds of either 0 or 1. Model performance was evaluated using Matthews correlation coefficient (MCC. Repeated five-fold stratified cross-validation with ten repeats was used to robustly estimate model performance. Cross-validation metrics were compiled and visualized using Python libraries Pandas, Matplotlib, and Seaborn, with MCC distributions presented as boxplots with overlaid strip plots.

The final druggability threshold was selected based on achieving the highest average MCC across evaluated thresholds. The full training dataset was used to train the final CatBoost model at the threshold that optimized MCC performance. This supervised training process is visualized in Fig 1d and implemented in the accompanying “stage2.py” file.

### Pocket druggability prediction and filtering

SiteMap pockets from candidate targets were filtered based on AlphaFold structure quality metrics. Pockets were retained only if both the pocket residues and the entire region spanning from the first to last amino acid in the pocket had at least 50% of amino acids with “very high” predicted AlphaFold quality (>90 pLDDT). Druggability predictions were then made for the quality-filtered pockets using the trained CatBoost model. Pockets were classified as druggable if the model predicted a probability greater than or equal to 50%. The prediction and filtering process is implemented in the “stage2.py” file and the process is visualized in Fig 1d.

### Pocket literature review

A manual literature review was conducted using Google Scholar and UniProt to identify representative natural and synthetic binders of each primary protein of interest, based on prior structural studies. To contextualize the location of these interactions, the residues involved in binding—whether natural or synthetic—were manually annotated and visualized using UCSF Chimera v1.18 (Pettersen et al., 2004) on the AlphaFold-predicted structure of the protein. Distinct sets of residues corresponding to each binder type were colored separately to enable visual differentiation of their respective binding regions. Only the residues on the parent protein structure were displayed; partner molecules were not included. These mapped residues were subsequently used to determine spatial overlap with predicted pockets during the contextual structural filtering step. Unstructured coils near the N-terminus were omitted from visualizations for clarity.

### Contextual structural analysis

To further filter and prioritize candidate binding sites for small-molecule intervention, we required that predicted SiteMap pockets overlap with both (1) the binding interface of a clinically validated antibody and (2) the site of a natural binder (e.g., a physiologically interacting protein partner). Only sites meeting both criteria were retained. For the resulting case, we selected the most recent high-resolution X-ray structure of the relevant protein–binder complex and applied InterProSurf to quantify the change in solvent-accessible surface area (ΔSASA) upon complex formation (Negi et al., 2007).

Residues were labeled as high-priority contact points if they (1) fell within the region overlapping both interfaces and (2) exhibited a ΔSASA greater than 50 Å^2^. These were interpreted as structurally validated interaction features that may inform small-molecule design aimed at mimicking therapeutically relevant contacts. ΔSASA values were visualized on the surface of the protein target of interest using ChimeraX 1.9, with a black-to-white gradient ranging from 0 Å^2^to the highest per-residue ΔSASA observed on the target protein (excluding binding partners).

### Static protein solvent mapping

Each target that passed the pocket literature review was analyzed using FTMap in protein–protein interface mode to identify putative druggable hotspots (Kozakov et al., 2015). AlphaFold-predicted monomeric structures were used as the primary input. In cases where a known post-translational modification (PTM) was located near a predicted binding site, the most recent available monomeric X-ray structure containing a representative PTM was also analyzed using FTMap to qualitatively assess whether the modification altered the local binding environment. This step was included as a precautionary measure, given that AlphaFold predictions may implicitly incorporate structural information from PTM-containing templates. FTMap output was downloaded and analyzed locally. A pocket was considered to show evidence of small-molecule druggability if at least one hotspot contained a minimum of 16 distinct probe types and was accompanied by one or two nearby secondary hotspots (Kozakov et al., 2011). Druggable hotspots were visualized in PyMol v2.5.4 with rainbow ribbon coloring, and unstructured N-terminal coils were hidden from all visualizations (Schrödinger, 2021).

### Dynamic protein fragment mapping

We utilized a SLURM cluster on Google Cloud Platform with preemptible compute nodes in conjunction with the SilcsBio Small Molecule Suite (2022.1 release) to calculate both regular and halogen-based SILCS FragMaps of AlphaFold structural predictions for targets that passed the druggability criteria from static protein solvent mapping (Ustach et al., 2019; SchedMD, 2022). FragMaps were visualized in PyMol, rainbow ribbon coloring was used, and unstructured coils near the n-termini were hidden. FTMap results and FragMaps were aligned in PyMol and FragMap clusters were labeled if within 1 angstrom of any of the FTMap probes.

### SILCS Hotspots mapping

To complement FTMap-based static solvent mapping, we used SILCS-Hotspots to evaluate potential fragment binding regions on the AlphaFold-predicted structure of each target (MacKerell et al., 2020). For targets with a known post-translational modification (PTM) near a binding site of interest, the most recent monomeric experimental structure containing a representative PTM was also analyzed to qualitatively assess whether the modification influenced predicted hotspot locations. All SILCS-Hotspots calculations were performed using the 2022.1 release of the SilcsBio Small Molecule Suite. The resulting hotspots were aligned with FTMap probe outputs in PyMol. Rainbow ribbon coloring was used and unstructured coils near n-termini were hidden. A SILCS hotspot was labeled and visualized if it fell within 5 Å of an FTMap probe molecule, based on results from either the AlphaFold or PTM-inclusive FTMap simulation. Hotspots meeting this criterion within a predicted binding site of interest were retained for further analysis and figure generation.

### Custom helper scripts

We used biopython, numpy, pandas, and seaborn libraries for data processing and basic analysis in our internal scripts (Cock et al., 2009; McKinney, 2010; Harris et al., 2020; Waskom, 2021).

## Results

### Model performance and target prioritization

Cross-validation of the CatBoost classifier demonstrated predictive performance across both binary druggability thresholding strategies. Classification performance was higher when distinguishing druggable and difficult pockets (labels = 1 or 2) from undruggable pockets (label = 0) using a threshold of > 0, compared to distinguishing druggable pockets (label = 2) from all others using a threshold of > 1, as measured by median Matthews correlation coefficient (MCC) values (Fig. 2a). The final model used for downstream prediction was trained at the threshold > 0, which yielded the highest mean MCC across cross-validation folds.

**Figure 2.**
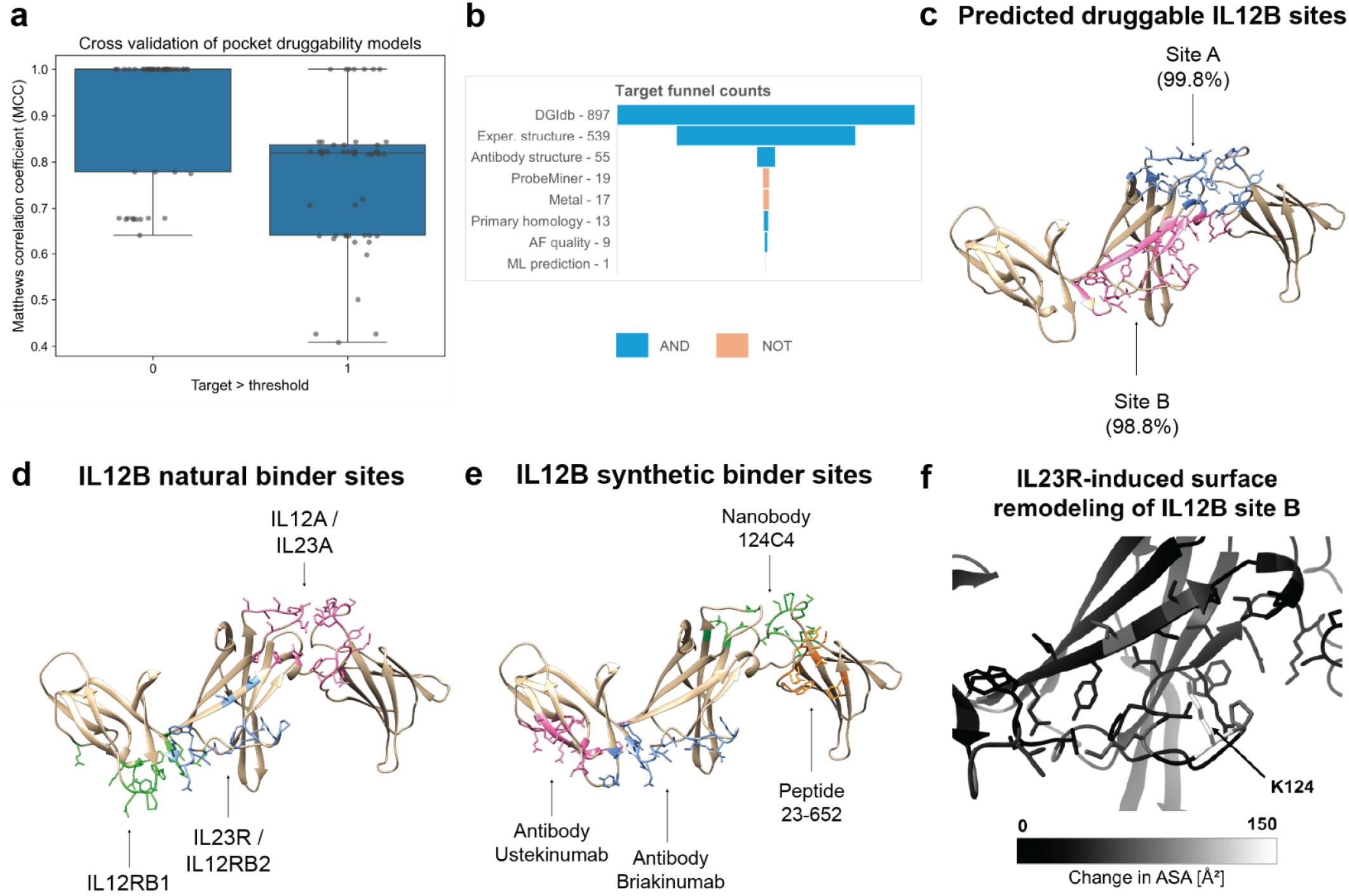
(a) 1/™1 of MCC corresponds to perfect agreement/disagreement with test data. Threshold interpretation in Fig 1d. (b) AF is AlphaFold and ML is machine learning. (c) Trimmed AlphaFold model; numbers in parentheses indicate CatBoost-predicted druggability confidence scores from the model with the highest mean MCC. (d) Trimmed AlphaFold model. (e) Trimmed AlphaFold model. Overlapping residues between ustekinumab and briakinumab antibodies not colored. (f) Data from InterProSurf using PDB 6WDQ structure; labeled residue has ΔSASA > 50 and overlaps with the briakinumab binding site.

The full target selection pipeline applied sequential filtering steps to candidate proteins: (1) identification as an approved or investigational target in DGIdb; (2) availability of at least one experimental structure in UniProt; (3) availability of at least one antibody-bound structure in the Thera-SAbDab database; (4) absence of existing small molecule ligands in the ProbeMiner database; (5) exclusion of proteins annotated with bound metal ions; (6) absence of significant primary sequence homology to other human proteins; (7) sufficient AlphaFold model quality in predicted pockets; and (8) predicted druggability based on the trained machine learning model. IL12B was the only target that satisfied all criteria (Fig. 2b).

### Site-level pocket identification in IL12B

Two predicted druggable pockets were identified in the IL12B AlphaFold structure. The CatBoost model predicted a 99.8% probability of druggability for Site A and a 98.8% probability for Site B. Site A included residues Y136, R160, Q166, E195, S197, A198, C199, P200, A201, A202, E203, E204, S205, L206, I229, R230, Y268, F269, D312, Y314, and Y315. Site B included H105, I111, W112, S113, T114, P123, K124, K126, T127, F128, L129, R130, C131, E132, W143, E221, N222, Y223, T224, S225, S226, F227, D231, I232, K234, S317, and S318 (Fig. 2c).

### Literature-informed structural binder mapping

A literature-based structural review was conducted to identify known natural and synthetic binders of IL12B. Residues involved in natural binding were identified for IL12RB1, IL23R/IL12RB2, and IL12A/IL23A based on prior structural studies (Yoon et al., 2000; Bloch et al., 2018, 2024; Glassman et al., 2021; Chen et al., 2024). Residues for synthetic binders were identified for ustekinumab, briakinumab, nanobody 124C4, and peptide 23-652 (Luo et al., 2010; Desmyter et al., 2017; Bloch et al., 2018; Pandya et al., 2020). The mapped residues for both natural and synthetic binders were visualized on the AlphaFold structure of IL12B (Fig. 2d–e).

### Receptor-induced remodeling of site B

The IL12B:IL23R heterodimer structure (PDB 6WDQ) was analyzed using InterProSurf to evaluate changes in solvent-accessible surface area (ASA) upon complex formation. At site B, residue K124 showed a ΔASA of 113.47 Å^2^ (Fig. 2f).

### Fragment-based solvent mapping of site B

FTMap simulations were conducted using both the AlphaFold model of IL12B (Fig. 3a) and an aligned crystal structure (PDB 1F42) that includes glycosylation near site B (Fig. 3b). In both cases, site B contained a primary cross cluster with at least 16 distinct probe types and additional nearby cross clusters.

**Figure 3.**
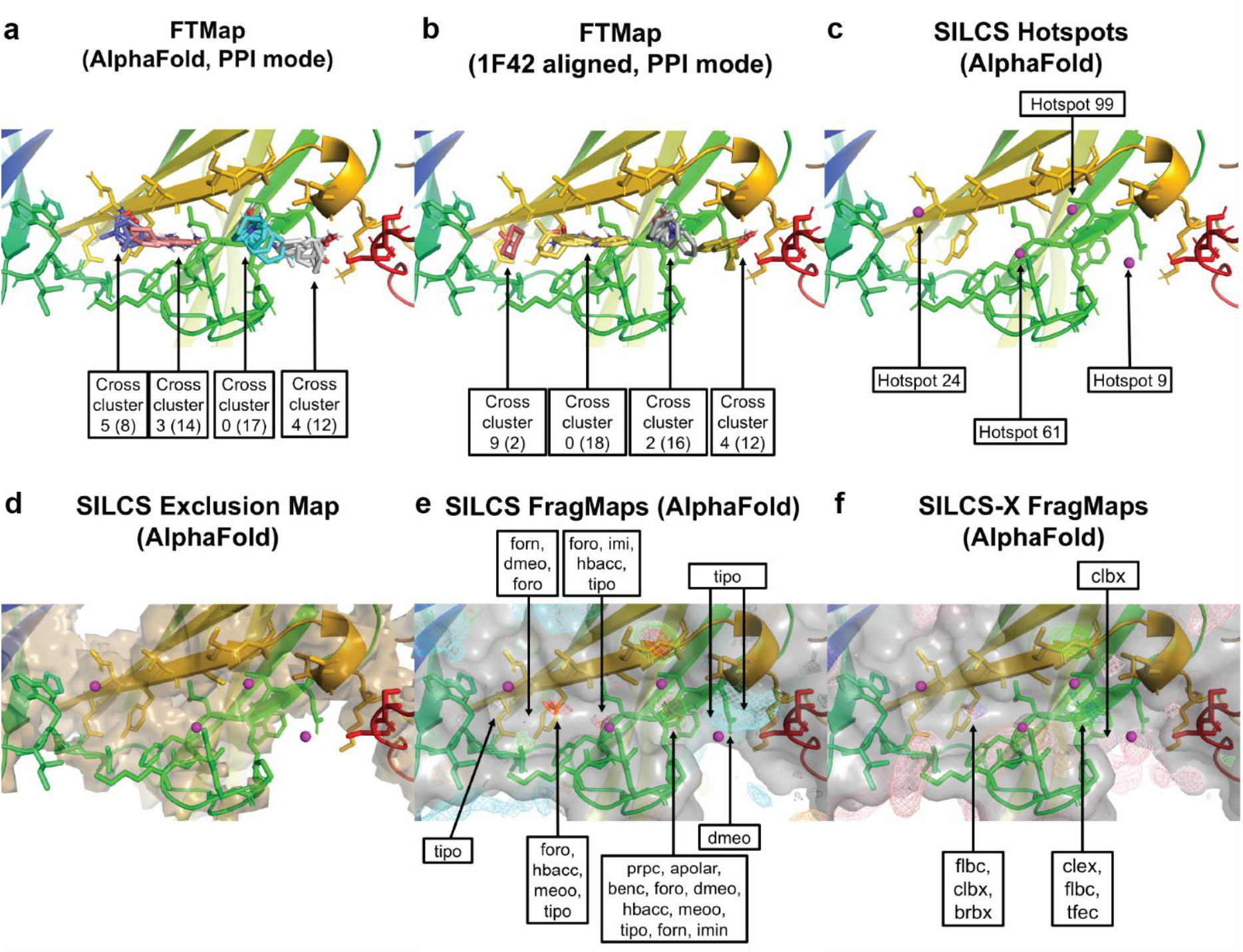
– Static and dynamic protein simulations of IL12B site B. Standard rainbow coloring of IL12B is used, and site B residues are shown in all subpanels. Reference labels for cross clusters (from FTMap static simulations) and hotspots (from SILCS dynamic simulations) reflect their original assignments from the respective raw simulation outputs. Numbers in parentheses for FTMap simulations are number of probes in cross cluster. AlphaFold-based models did not incorporate any post-translational modifications. (b) The 1F42-based results derive from an experimental IL12B structure that includes glycosylation near site B; cross clusters are aligned to and visualized on the AlphaFold IL12B structure for consistency. (e–f) SILCS and SILCS-X probe identities and physicochemical properties are summarized in Table 1.

SILCS-Hotspots analysis identified multiple predicted fragment-binding regions at site B, including hotspots 9, 24, 61, and 99 (Fig. 3c). Hotspot 99 was partially buried, as shown in the SILCS exclusion map (Fig. 3d), and located within 10 Å of other hotspots. Hotspot 99 was nearby one apolar FragMap region and a nearby hydrogen bond acceptor (hbacc) region. Additional hbacc density was observed between hotspots 24 and 61 and near hotspot 61 (Fig. 3e). SILCS-X FragMaps showed significant spatial overlap with corresponding SILCS FragMaps at site B (Fig. 3f).

**Table 1.**
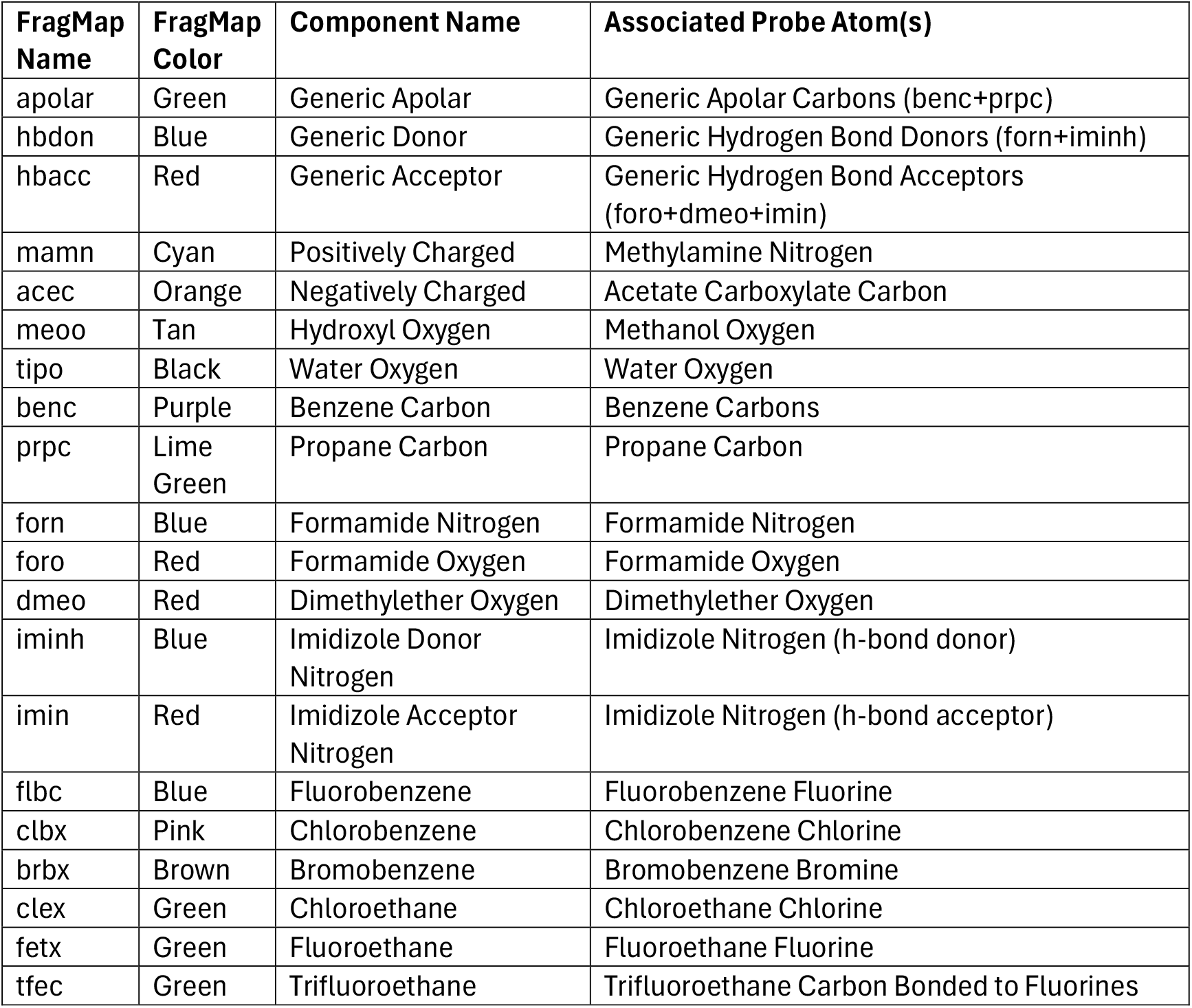
SILCS and SILCS-X FragMap probe reference. Probe names, colors, component types, and key interaction atoms used in standard SILCS and halogen-augmented SILCS-X FragMap simulations. Standard probes include neutral, polar, and charged fragments; halogen probes represent specific halogenated functional groups. Colors refer to visualizations used throughout figures in this study.

## Discussion

The multi-parameter prioritisation workflow reduced an initial DGIdb set of 897 drug-associated genes to a single high-confidence target, IL12B, and highlighted two predicted pockets. Site B drew particular interest because it overlaps part of the briakinumab epitope and has no reported small-molecule ligands.

Static protein solvent mapping with FTMap provided consistent evidence that site B can sustain fragment binding. The largest cross cluster contained 17 probe types in the AlphaFold model and 18 in the experimentally derived, glycosylated structure; in both cases additional nearby cross clusters were present. This pattern exceeds the ≥16-probe threshold, with supporting clusters, that has been correlated with druggability in benchmark studies (Kozakov et al., 2011).

Dynamic fragment mapping with SILCS-Hotspots corroborated the location of the pocket by identifying at least one partially buried hotspot with neighboring hotspots within <10 Å. Although only one apolar FragMap region was detected, the co-localised hydrogen-bond acceptor density fulfils other SILCS prioritisation criteria and indicates chemical features that could guide fragment elaboration.

Clinically, IL12B is already validated by monoclonal antibodies such as ustekinumab and briakinumab, which are effective in inflammatory bowel disease (IBD) and psoriasis (Al-Bawardy et al., 2021). Antibody therapy, however, entails parenteral dosing and prolonged systemic exposure that complicate infection management; current IBD biologics have been linked to markedly increased infection rates, leading treatment guidelines to recommend temporary withdrawal during selected infections (Zabana et al., 2019; Kucharzik et al., 2021). A small-molecule inhibitor directed to site B could, in principle, offer oral delivery, shorter half-life, and improved mucosal penetration—attributes likely to facilitate rapid dose modulation and reduce systemic immunosuppression.

Taken together, the convergent FTMap and SILCS findings nominate site B as a previously unexploited pocket for small molecule development on IL12B. Experimental characterisation of this site, including fragment screening and assessment of conformational and post-translational heterogeneity, now represents a logical next step toward broadening the therapeutic arsenal beyond existing antibody modalities.

### Analytic limitations

This study is limited by its reliance on computational tools and simulations, without direct experimental validation. The machine learning model used for druggability prediction was trained on a relatively small and class-imbalanced dataset, which may affect its generalizability to uncharacterized targets.

A central limitation of this analysis involves uncertainty regarding the physiological structural state of IL12B. While IL12B can exist as a monomer, homodimer, or as part of heterodimers with IL12A or IL23A, the structural and functional relevance of each state may vary by context. The AlphaFold structural model used in this study reflects a monomeric configuration, which may not fully represent the biologically active conformations encountered in vivo.

Additionally, the post-translational modification (PTM) status of IL12B remains incompletely characterized. Glycosylation patterns, in particular, may differ across subcellular compartments (e.g., intracellular vs. extracellular) and may be influenced by tissue type, inflammatory state, or disease context. Structural models used in this study do not account for potential variety in PTMs, and they may affect both the geometry and accessibility of predicted small-molecule binding sites. The 1F42 crystal structure, one of the few experimental structures for monomeric human IL12B, lacks accompanying electron density maps and dates from before modern structure validation standards, limiting its utility for refinement. Further experimental work is needed to define the in vivo conformational landscape of IL12B in human patients, including its tissue-specific PTM states and physiologically dominant oligomeric forms.

## Conclusion

Our integrated in silico workflow—combining stringent sequence, structural, and machine-learning filters with complementary solvent-mapping techniques—surfaced site B on IL12B as a tractable yet chemically unexplored pocket. FTMap clustering and SILCS-Hotspots converged on this region, together indicating a permissive blend of polar and apolar interaction motifs that should accommodate fragment growth and, ultimately, lead optimisation. Critically, site B lies outside the canonical dimer interface yet overlaps part of the briakinumab epitope, anchoring its biological relevance while remaining untouched by small-molecule medicinal chemistry.

Translating these findings could expand the therapeutic repertoire for inflammatory bowel disease and related disorders, where antibody therapies are effective but hampered by parenteral dosing and prolonged systemic exposure. A well-designed oral inhibitor of IL12B has the potential to deliver adjustable pharmacokinetics, improved mucosal penetration, and rapid dose modulation—attributes that align with clinical needs in infection-prone patient populations.

The next milestones are clear: obtain high-resolution structures of IL12B in its predominant PTM and oligomeric states, initiate fragment screening against site B, and profile early hit series for selectivity. Success on these fronts would open a direct path toward the first small-molecule IL12B modulator, offering industry and academia a fresh entry point into a validated yet under-leveraged cytokine target.

## Acknowledgements

We gratefully acknowledge the support of several organizations that made this work possible. We thank the Google for Startups Cloud Program for generously providing $100,000 of computational credits to Epilog, L.L.C. We are also appreciative of Schrödinger, L.L.C. and SilcsBio, L.L.C. for providing trials of their software packages. Chemaxon generously provided a complimentary license for their software (Chemaxon, 2024). The Chemicalize platform was usefulf for visualizing chemical structures documented in various crystallization protocols for the IL12B protein extracted from the literature. We also acknowledge Wingware for access to its Wing Pro IDE for Python development (Wingware, 2025).

Salt Lake Community College (SLCC) played a crucial role in the development of this research through two key programs. Within SLCC’s STEM Learning department, Dr. Lane Law and Dr. Santosh Kiran Balijepalli offered invaluable guidance through their STEM 2010 class. Their expert feedback significantly enhanced the author’s ability to articulate, refine, and communicate scientific ideas. The Everyday Entrepreneur Program, taught by Jon Beutler, provided foundational training in market research, competitive analysis, and minimum viable product development.

The author also thanks Drs. Peter Iles and Tim Beagley for sparking his interest in the life sciences through engaging introductory and cell biology courses, and Dr. Craig Thulin for strengthening his understanding of biochemical principles.

The foundational skill sets used in this work were first developed during two summers the author spent in the National Science Foundation’s Research Experience for Undergraduates (REU) program (NSF award #0835713) at Harvard, working in the Aspuru-Guzik group under the mentorship of Dr. Dmitrij Rappoport. The author is especially grateful to Dr. Rappoport for his generous mentorship and for offering thoughtful feedback that helped improve this manuscript.

Finally, we acknowledge the use of advanced language models, including Claude 4 Sonnet (Anthropic) and ChatGPT (OpenAI), which assisted with editing titles, improving abstract clarity, and refining the presentation of technical content.

## Conflict of interest disclosure

Trevor J. Tanner, the founder of Epilog, L.L.C., received computational resources through the Google Cloud for Startups program. Some of these resources, along with academic resources and affiliations, contributed to the research process used in the identification of the IL12B target described in this work. Epilog, L.L.C. asserts no rights associated with any of the findings presented herein. The author declares no financial conflicts of interest related to this work.

### Data, scripts, and code availability

Data, scripts, and code are available online: https://doi.org/10.5281/zenodo.15765054. Some intermediate results generated using Schrödinger’s commercial software have been excluded from the public repository to comply with Schrödinger’s End User License Agreement. Certain scripts may not function without Schrödinger-specific outputs and are provided for reference. Recreating the original dataset may require a Schrödinger software license. All other non-Schrödinger files necessary for understanding and reproducing our work are included. The repository README provides detailed instructions.

